# Can a fish learn to ride a bicycle? Sensorimotor adaptation to destabilizing dynamics in the weakly electric fish *Eigenmannia virescens*

**DOI:** 10.1101/2023.01.27.525956

**Authors:** Yu Yang, Dominic G. Yared, Noah J. Cowan

## Abstract

Humans and other animals can readily learn to compensate for destabilizing dynamics, such as balancing an object or riding a bicycle. How does the nervous system learn to compensate for such destabilizing dynamics, and what are the benefits of the newly learned control policies? To investigate these questions, we examined how the weakly electric glass knifefish, *Eigenmannia virescens*, retunes its control system in the face of novel, destabilizing dynamics. Using a real-time feedback system, we measured swimming movements as seven individual fish tracked a moving refuge, and we fed the swimming movements back through novel dynamics to alter the refuge motion, creating an artificially destabilizing reafferent loop. We discovered that fish learned to retune their sensorimotor controllers as the artificially destabilizing feedback was gradually introduced. Furthermore, when the artificial feedback was extinguished, fish exhibited a clear aftereffect, retaining their learned sensorimotor controllers for several minutes before washing out. This retuning of the control system under destabilizing dynamics: (i) improved tracking performance compared to the predicted performance had fish not re-tuned their baseline controller, (ii) reduced sensitivity of the sensorimotor system to low-frequency disturbances, such as would arise from turbulence or motor noise, and (iii) improved phase margin, a measure of stability robustness, despite the artificial feedback driving the putative baseline control system towards instability. Our study sheds light on how the nervous system adapts to changing closed-loop dynamics, and how those changes impact performance and stability; the presence of aftereffects suggest a plasticity-based mechanism reminiscent of cerebellar learning.

## 2 Introduction

Animals routinely alter their behavior in response to novel sensorimotor feedback. For example, insects can adjust to limb amputation [1–4], antenna trimming [5, 6], and wing damage [7–9] caused by predation or collisions, and in some cases, they may make these adjustments without learning [2]. Of course, humans and other animals can adeptly learn new motor behaviors and tune existing ones through practice [10], and the mechanisms and strategies for such learning are being revealed [10], including at the circuit level [11–13]. However, from the perspective of control theory, it is unclear how animals learn novel dynamics, and how the updated controllers impact system level performance. Do animals simply maintain stability [14], or do they improve other aspects of performance? Do they favor robustness [15] or optimality [16]? To address these questions, we studied a well-suited model organism, namely the weakly electric glass knifefish *Eigenmannia virescens*, and considered its refuge tracking system as a control loop; we identified how the fish’s sensorimotor controller changed under artificial, destabilizing feedback.

*Eigenmannia virescens* has been likened to an “aquatic hummingbird” [17], smoothly hovering in place or making agile movements to track a moving refuge [18]. These fish swim back and forth to sense their position within such a refuge [19–21], while also tracking the movements of the refuge [18,22,23]. These fore-aft swimming movements are produced by thrust forces generated by adjustments to the counter-propagating waves of its long, ventral ribbon fin [15,24]. *Eigenmannia* also relies on a robust feedback control system that is insensitive to the locomotor (plant) dynamics [15], and thus may not need to adapt to morphophysiological variability [15].

Experiments to understand how learning occurs often involves artificially altering sensorimotor feedback, like walking on split-belt treadmills [25, 26], visuomotor rotations and mirror reversal while tracking moving targets [27, 28], and reaching for targets under artificially induced force-fields [29, 30]. In our present study, we examined sensorimotor adaptation in fish refuge tracking by designing an artificial feedback loop that could be used to alter the closed-loop dynamics. This experimental manipulation is analogous to inserting a dynamical system between the rotation of a steering wheel, and the turning of the front wheels of a drive-by-wire automobile; in this way, the steering of the car could become non-intuitive and even unstable, if the driver failed to adapt their sensorimotor controller. To destabilize the closed-loop refuge tracking system, we increased the gain of the artificial feedback, systematically measuring the sensorimotor controller of each fish using system identification (i.e., stimulus–response modeling). Finally we removed the artificial feedback to assess any aftereffects in the sensorimotor controller, as an indicator of learning.

## 3 Significance

Learning to adapt to new system dynamics—like balancing a pole on your finger, riding a bike, or learning to walk—requires making changes to the complex transformation between sensory percepts and muscle commands. Here, we take a comparative approach to understand how the nervous system changes its controller in the face of new dynamics, how this learning improves task performance, and show that the controller is learned through a plasticity mechanism. This research suggests that neural substrates for learning to control complex, destabilizing locomotor dynamics—like riding a bicycle—are shared across vertebrate taxa.

## 4 Results

### 4.1 Fish tuned their controller dynamics in response to destabilizing feedback

We designed an artificial feedback loop to alter the closed-loop dynamics that fish experienced during refuge tracking (Figs. 1(A, B)). Specifically, we fed fish position back through a high-pass filter in real time and added it to a pre-defined pseudo-random sum-of-sines input (Fig. S1(A)) [23, 28]. To drive the closed-loop system towards instability, we incrementally increased the artificial feedback gain, *k*, on the high-pass filter (Fig. 1(C)). The experiments started from open-loop “baseline,” followed by a “closed-loop” period in which the high-pass feedback gain was gradually increased in discrete steps. This feedback was designed based on preliminary analysis [31] to destabilize the entire system in an effort to elicit learning. Finally, we removed the artificial feedback to assess aftereffects (see Materials and Methods, section 9.3).

**Figure 1.**
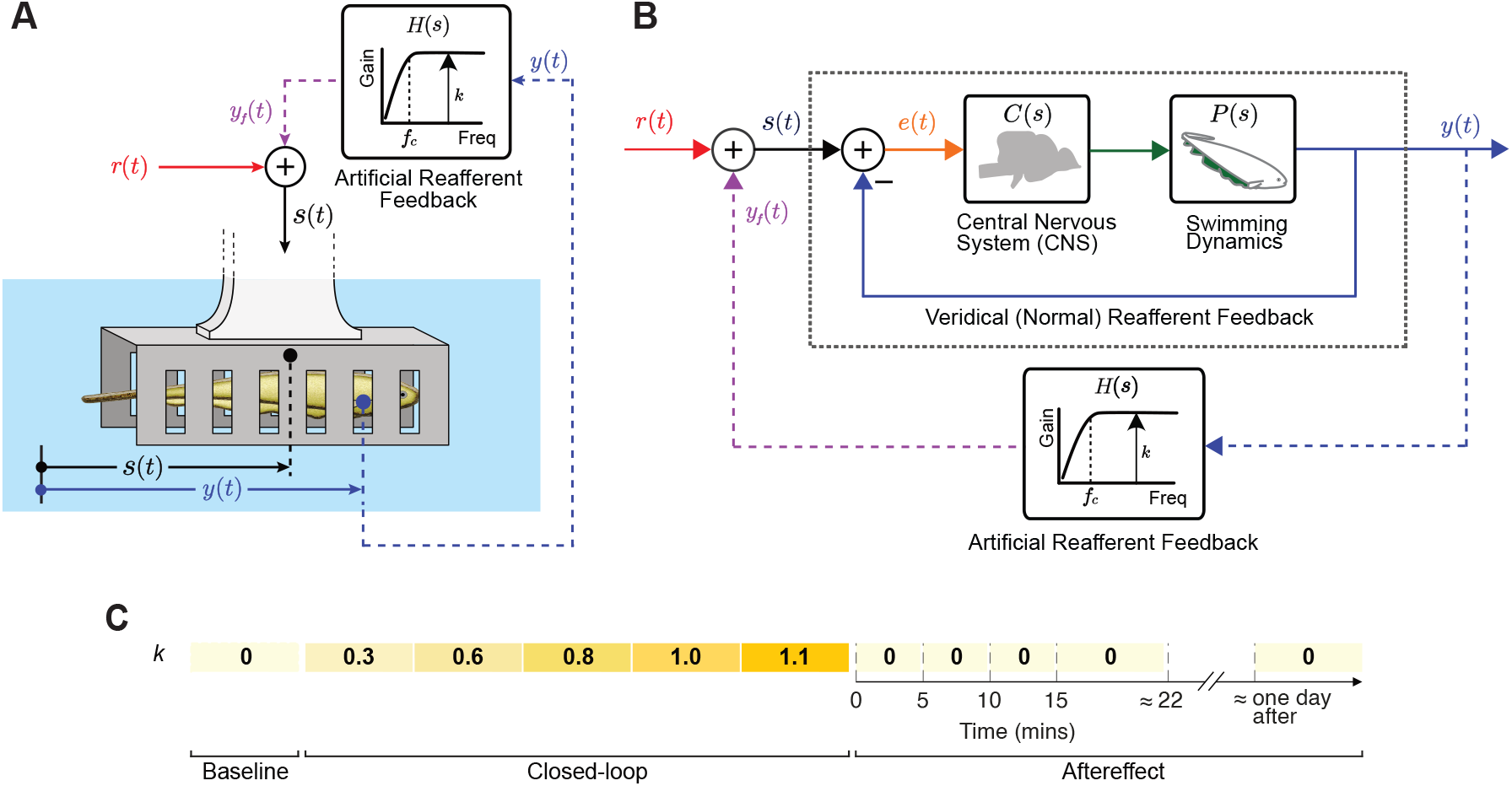
Experimental protocol and stimulus design. (A) Schematic of artificial, destabilizing feedback system. The longitudinal fish position, *y*(*t*), was measured in real time using a custom video tracking system, and processed through a high-pass filter, *H*(*s*) with gain *k* and cut-off frequency *f_c_* = 0.16 Hz. The high-pass-filtered position, *y_f_* (*t*), was then added to a pseudo-random (sum-of-sines) reference signal *r*(*t*), and fed to the refuge, i.e., *s*(*t*) = *y_f_* (*t*) + *r*(*t*), creating an artificial reafferent loop. (B) Block diagram of the closed-loop system depicted in (A). The artificial, high-pass filter feedback is closed around the fish sensorimotor system (black dotted box). The fish is presumed to sense the difference between the longitudinal refuge position *s*(*t*), and its own reafferent feedback *y*(*t*). This difference *e*(*t*) (“sensory slip” [19]) is then processed by the nervous system, *C*(*s*), and swimming dynamics, *P* (*s*). (C) Protocol: Experiments started from an experimental open-loop “baseline” period (*k* = 0), followed by “closed-loop” period in which we incrementally increased the artificial high-pass filter gain *k* from 0.3 to 1.1 over a course of about 33 minutes. Finally, we extinguished the artificial feedback to test for an “aftereffect” (*k* = 0 again). The sensorimotor controller was tested at five time intervals during the “aftereffect” period: 0 to <5 minutes, 5 to <10 minutes, 10 to <15 minutes, 15 to around 22 minutes, and approximately one day after the end of the “closed-loop” period.

To uncover how the artificial feedback changed the fish’s tracking response, we performed frequency domain system identification on seven individual fish (Materials and Methods) to estimate the empirical frequency response functions of individual fish. The frequency response functions we estimated comprised the fish’s own reafferent feedback (see black dotted box in Fig. 1(B)), but excluded the artificial feedback. Using this method, we assessed how changes to the artificial feedback (via the gain *k*) led to changes in the fish’s control system (as measured through its frequency response).

We found that fish systematically tuned their controllers under the artificial feedback. Fish added phase lead at intermediate frequencies (around 0.65 Hz) and increased sensorimotor gain, especially at higher frequencies (above 1 Hz), in response to the increasing artificial feedback gain *k* (Fig. 2(A)). This trend of increased gain and phase was seen in each of the seven fish (Figs. 2(B, C)).

**Figure 2.**
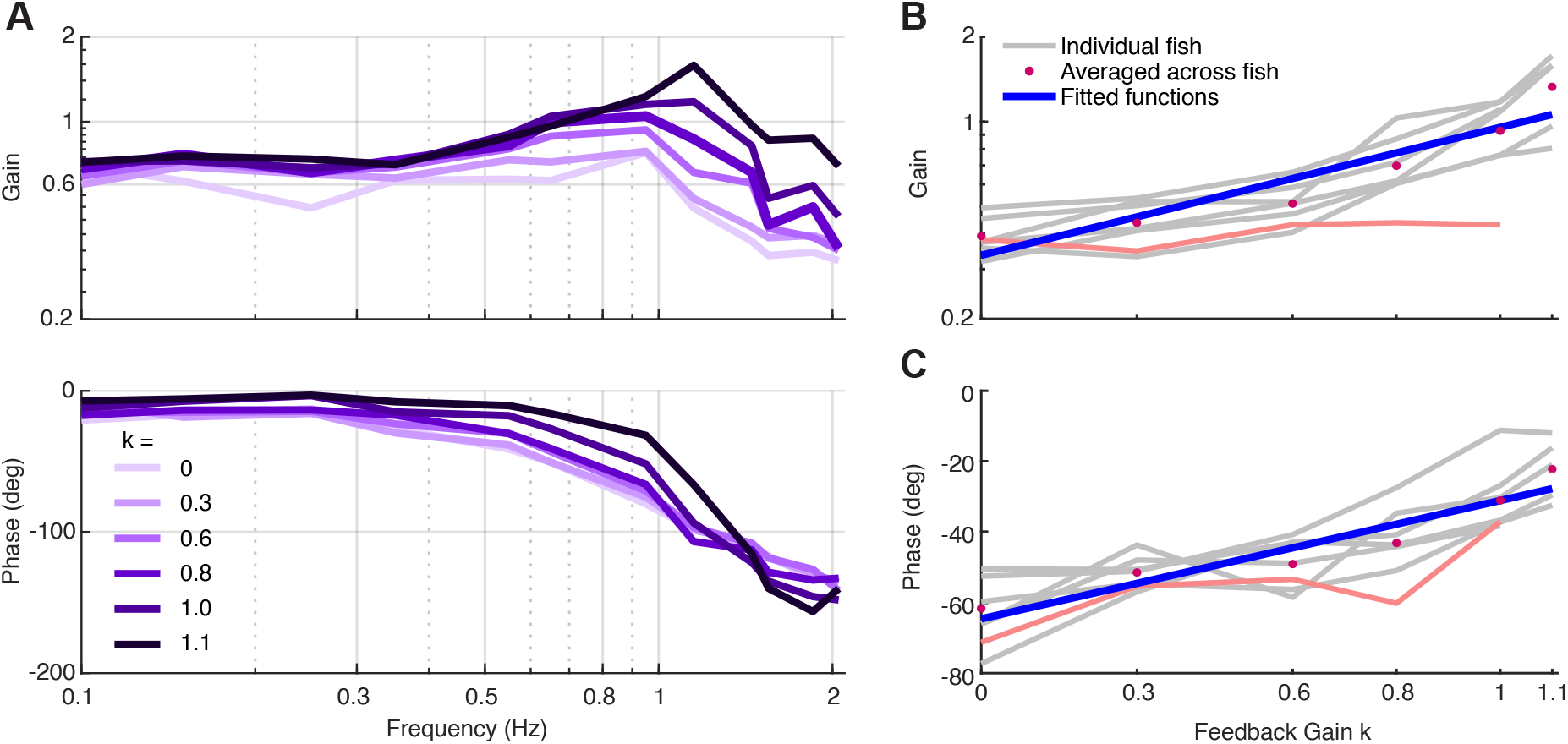
Fish tuned their controller dynamics in response to destabilizing feedback. (A) Frequency response of one representative fish. As the feedback gain was increased, the gain of fish frequency response function also increased (especially at frequencies *>* 0.95 Hz) and the phase lag decreased. (B) Gain of fish frequency response functions at 1.15 Hz and (C) phase at 0.65 Hz as a function of feedback gain *k* (seven fish for most points, except for *k* = 1.1 where two fish was excluded due to poor data quality). Linear functions (blue lines) were fitted to the averaged gains (pink dots) in logarithmic scale at 1.15 Hz, and averaged phases (pink dots) in linear scale at 0.65 Hz across all individual fish (gray curve). Six out of seven fish exhibited significant (*p <* 0.05) positive correlations between high-pass gain *k* and log of gain at 1.15 Hz (*ρ >* 0.91) or phase (*ρ >* 0.86) at 0.65 Hz; one fish (red line) exhibited positive, but non-significant correlations for both gain (*ρ* = 0.75) and phase (*ρ* = 0.8).

### 4.2 Fish temporarily retained their learned controller, exhibiting a clear after-effect

We tested if aftereffects persisted when the closed-loop feedback was abruptly removed, as the presence of such aftereffects would indicate a plasticity-based mechanism, reminiscent of cerebellar learning [25, 26, 28]. We averaged the frequency responses in blocks of three or five trials (details see Materials and Methods, section 9.3) during the “aftereffect” period (see Fig. 1(C)), and compared the result with the “baseline” frequency response. Fig. 3(A) shows the averaged fish frequency response during the “aftereffect” period, as well as the “baseline” period, from one representative fish. In each frequency response plot, the fish was under normal, i.e., veridical feedback, with no artificial high-pass filter in the feedback loop. Note that just after the artificial feedback was removed, both the frequency response gain and phases remained elevated, revealing that the tuned controller persisted right after the feedback was removed. Then, over 15 minutes, the frequency responses returned to baseline, as the learned controller was washed out.

**Figure 3.**
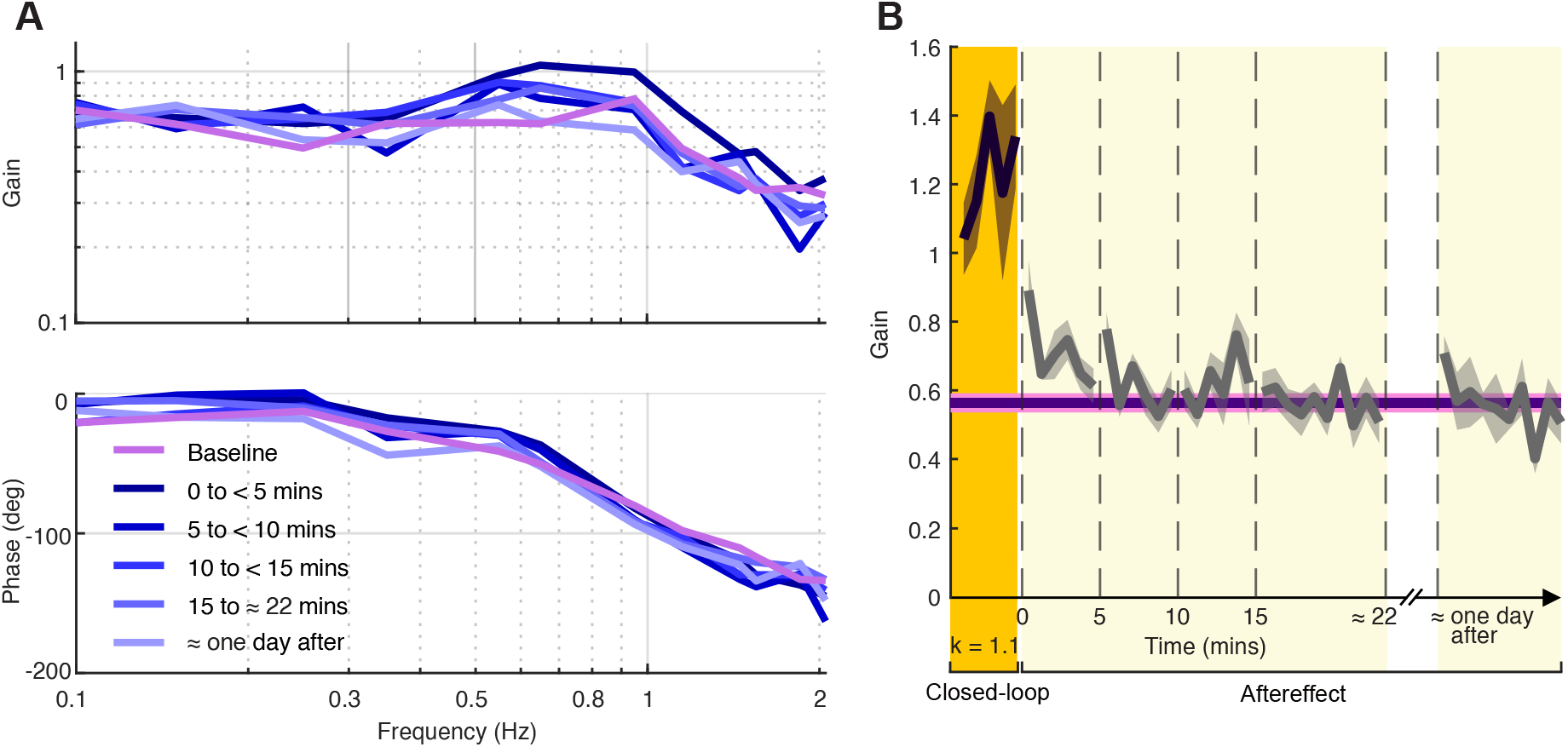
Fish temporarily retained learned transfer function in “aftereffect” period. (A) Averaged fish frequency response from cases: baseline, 0 to *<*5 minutes, 5 to *<*10 minutes, 10 to *<*15 minutes, 15 to around 22 minutes, and approximately one day after the end of closed-loop experiments from online tracking data for one fish. The frequency response gain and phases of the averaged “0 to *<*5 mins after” case remained elevated, and washed out within about 15 minutes. (B) Fish frequency response gain at 0.95 Hz from late learning “*k* = 1.1” (when *t <* 0), “0 to *<*5 mins after,” “5 to *<*10 mins after,” “10 to *<*15 mins after,” “15 to around 22 mins after,” and “approximately one day after” cases, by bootstrapping online tracking data (10, 000 times) across six fish. The gray solid lines are means and shaded regions are standard deviations (SDs). The purple solid line is the mean of bootstrapped data in “baseline,” while adjacent pink shaded region is SD.

To test whether this aftereffect was significant, we bootstrapped data across six fish (Fig. 3(B)), and found that the average frequency response gain at 0.95 Hz of the first 20 seconds of the “0 to <5 mins after” experimental period was markedly lower than the average gain in closed-loop (*k* = 1.1), but still remained significantly elevated above baseline. The frequency response gain then gradually decayed towards baseline within about 15 minutes. Similar trends were found in frequency response gain at 1.15 Hz and frequency response phase at 0.65 Hz (Fig. S2).

### 4.3 Advantage (i): tuned controller improved tracking performance

Fish systematically tuned their controllers under the artificial, destabilizing feedback. To understand the possible benefits of this adaptive control process, we examined how the tuned controllers impacted fish tracking performance. We computed the normalized error *|E*(*jω*)*/R*(*jω*)|, i.e., the magnitude of fish sensory slip *e*(*t*) over reference *r*(*t*) in the frequency domain, for all five feedback gains. We used the individual baseline controller for each fish, estimated without the artificial, high-pass-filter feedback, to predict what would happen in the theoretical case that the fish did *not* tune its controller when we introduced the destabilizing artificial feedback.

We indirectly verified that the fish tuned their controllers to mitigate errors in the tracking performance. We inferred this because had the fish *not* tuned their controllers, the destabilizing feedback would have worsened tracking performance, particularly below about 0.6 Hz in most fish (for example, see Fig. 4A, top). However, the predicted error was largely suppressed by the observed tuned controller (as shown for the same fish in Fig. 4A, bottom). To quantitatively compare the observed versus predicted normalized error, we integrated the normalized observed and predicted tracking error in the frequency domain for all seven fish. For all seven fish, and nearly all gains (32 of 33), the observed error for the tuned controller was significantly less than that predicted when assuming no tuning (*p <* 0.001, two tailed Mann-Whitney-Wilcoxon test; Fig. 4B). In summary, the tuned controllers decreased the tracking error compared to the prediction assuming no tuning.

**Figure 4.**
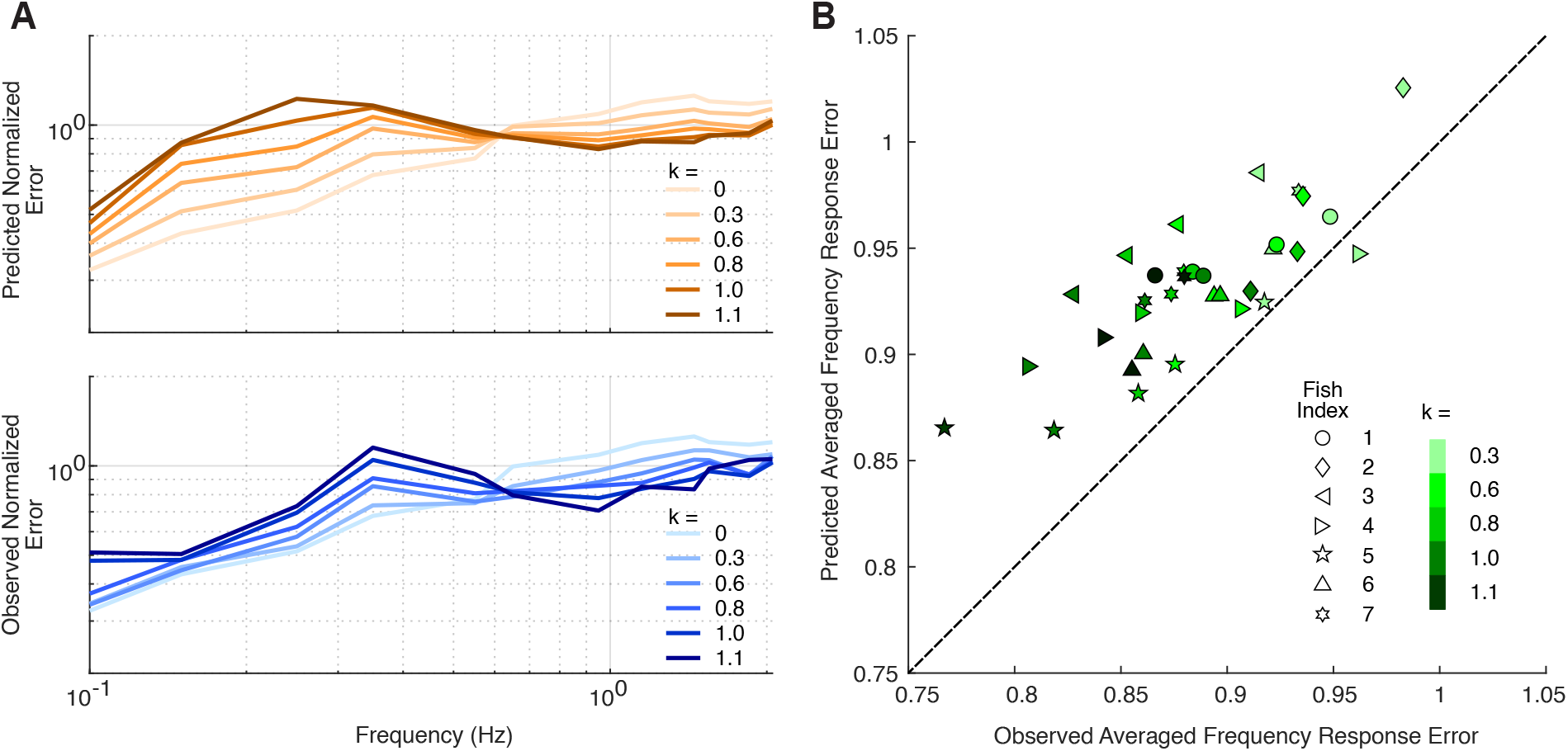
Tuned controller improved tracking performance. (A) The predicted normalized error *E*(*jω*)*/R*(*jω*) was computed by assuming that the fish did not tune their baseline controller as the feedback gain was increased; *R*(*jω*) and *E*(*jω*) are the frequency domain reference signal *r*(*t*) and “sensory slip” *e*(*t*), respectively. As can be seen for a representative fish (top), the predicted normalized error increased substantially below about 0.6 Hz, and decreased slightly for higher frequencies. In contrast, the observed error measured for the same fish (bottom) was substantially lower, as the tuned controller largely mitigated the effects of the destabilizing feedback. (B) Scatter plot showing the predicted and observed averaged frequency response error, integrated across the frequency range 0.1 2.05 Hz, for each feedback gain, for seven fish. A total of 32 of 33 points are located above the identity line, showing that the tuned controller decreased normalized tracking error over predicted normalized tracking error for all seven fish, and for nearly all gains (32 of 33) (*p <* 0.001, two tailed Mann-Whitney-Wilcoxon test).

### 4.4 Advantage (ii): tuned controller reduced sensitivity to disturbance

The *sensitivity function S* (see Materials and Methods) reflects how well a feedback system suppresses disturbances [32], such as motor noise [33], turbulence [34], and other external forces. At frequencies *ω* for which *|S*(*jω*)*| <* 1 (improved *robustness*), the system *attenuates* disturbances. Conversely, the system *amplifies* disturbances when *|S*(*jω*)*| >* 1 (increased *fragility*) [32, 35]. Critically, for many systems, there is a tradeoff between “robustness” and “fragility” because of a general result, known as the “waterbed effect” [32], which states that the integral of the natural log of the sensitivity function magnitude—i.e., the *Bode sensitivity integral* —is 0:

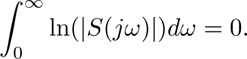

This result depends on technical details of system dynamics [32] (see SI Appendix, Supporting Information Text section 10.1.1 for details). This integral constraint produces a “conservation law,” implying that suppressing disturbances (ln *|S| <* 0, improving *robustness*) at some frequencies requires amplifying disturbances (ln *|S| >* 0, increasing *fragility*) at other frequencies.

We computed the predicted sensitivity function for each feedback gain, *k*, based on each fish’s baseline controller (estimated without the artificial, high-pass-filter feedback applied). This predicted sensitivity was compared to the sensitivity function that each fish achieved when the feedback was applied. If the fish had not changed their controller, then the predicted and observed sensitivity functions would have been identical. For both the predicted and observed sensitivity functions, we approximated the Bode integral over the experimental bandwidth of 0.1 *−* 2.05 Hz (0.63 *−* 12.9 rad/s) using trapezoidal integration. Fig. 5(A) shows the natural logarithm of sensitivity function magnitude (predicted and observed) at *k* = 1.1 from one representative fish over this bandwidth; the Bode integral was approximated as the difference of the positive and negative areas of this function. Fig. 5(B) shows that, across fish, as the feedback gain *k* was increased, the Bode integral became increasingly negative compared to the predicted integral (*p <* 0.05 for *k* = 0.3, *p <* 0.01 for each *k >* 0.3, paired-sample *t* test).

**Figure 5.**
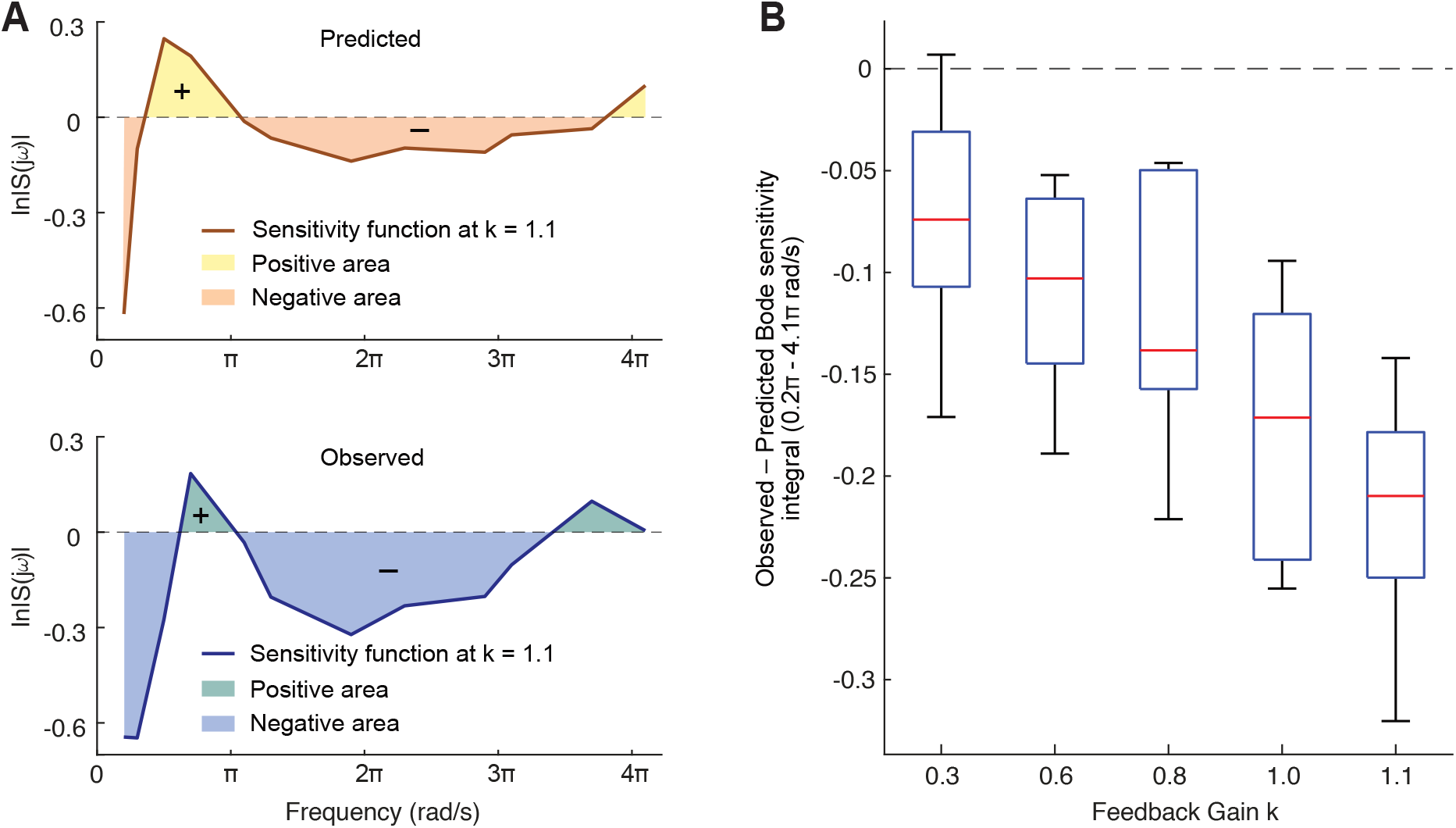
Tuned controller reduced sensitivity to disturbance. (A) Predicted (based on baseline controller) and observed (based on fish’s tuned controller) Bode sensitivity functions for *k* = 1.1, from one representative fish. The natural logarithmic magnitude of observed Bode sensitivity function has a larger negative region than predicted (See Fig. S3 for larger frequency range). (B) For each individual fish, and at each gain *k*, we computed the difference between the observed and predicted Bode sensitivity integrals over the frequency range 0.2*π*–4.1*π* rad/s. Box and Whisker plots of these values are shown for all seven fish for *k* = 0.3 1.0, five fish for *k* = 1.1. The mean value across all fish was *<* 0 for each gain (*p <* 0.05 for *k* = 0.3, *p <* 0.01 for the remaining gains, paired-sample *t* test for observed vs. predicted Bode sensitivity integral).

This finding indicates that the change of fish controller brought about another benefit, i.e., reducing sensitivity to disturbances within this relatively low frequency band (*<* 2.05 Hz). Moreover, this enhancement over prediction became more pronounced with ever-increasing artificial feedback gains, as the difference between the observed and predicted Bode integral decreased with increased gain *k* (*p <* 0.05, correlation coefficient *ρ* = *−*0.94 for the average across seven fish). Such tuned, low-frequency disturbance rejection may help suppress turbulence and other environmental and neuromuscular perturbations as we consider further in the discussion.

### 4.5 Advantage (iii): tuned controller improved phase margin

We examined how changes in the fish’s sensorimotor controller affected closed-loop stability. Similar to our previous analyses in this paper, we used the fish’s baseline frequency response—before the destabilizing artificial high-pass filter was introduced to the feedback loop—to mathematically predict closed-loop performance, and compared this baseline prediction to the observed tuned control system. We observed that fish maintained or improved measures of stability robustness, compared to what would be expected if they had not tuned their controller, as is now described.

To quantify stability robustness, we used two concepts from control theory, namely, the *Gain Margin* and *Phase Margin* which provide measures of how much a system can be perturbed before it becomes unstable. More precisely, *Gain Margin* (GM) measures how close the gain of the loop function is to unity gain (0 dB) at the frequency where the phase of the loop function crosses *−*180*^◦^*. Likewise, the *Phase Margin* (PM) measures how much the phase of the loop function is above *−*180*^◦^* at the frequencies where the gain crosses 0 dB (see Fig. 6A). Here, the loop function *L* = *CP* (1 *− DH*) includes the fish’s own controller and plant, *C*(*s*) and *P* (*s*), the system delay in the forward loop *D*(*s*) (see SI Appendix, Supporting Information Text section 10.1.3), as well as the artificial high-pass filter feedback, *H*(*s*).

**Figure 6.**
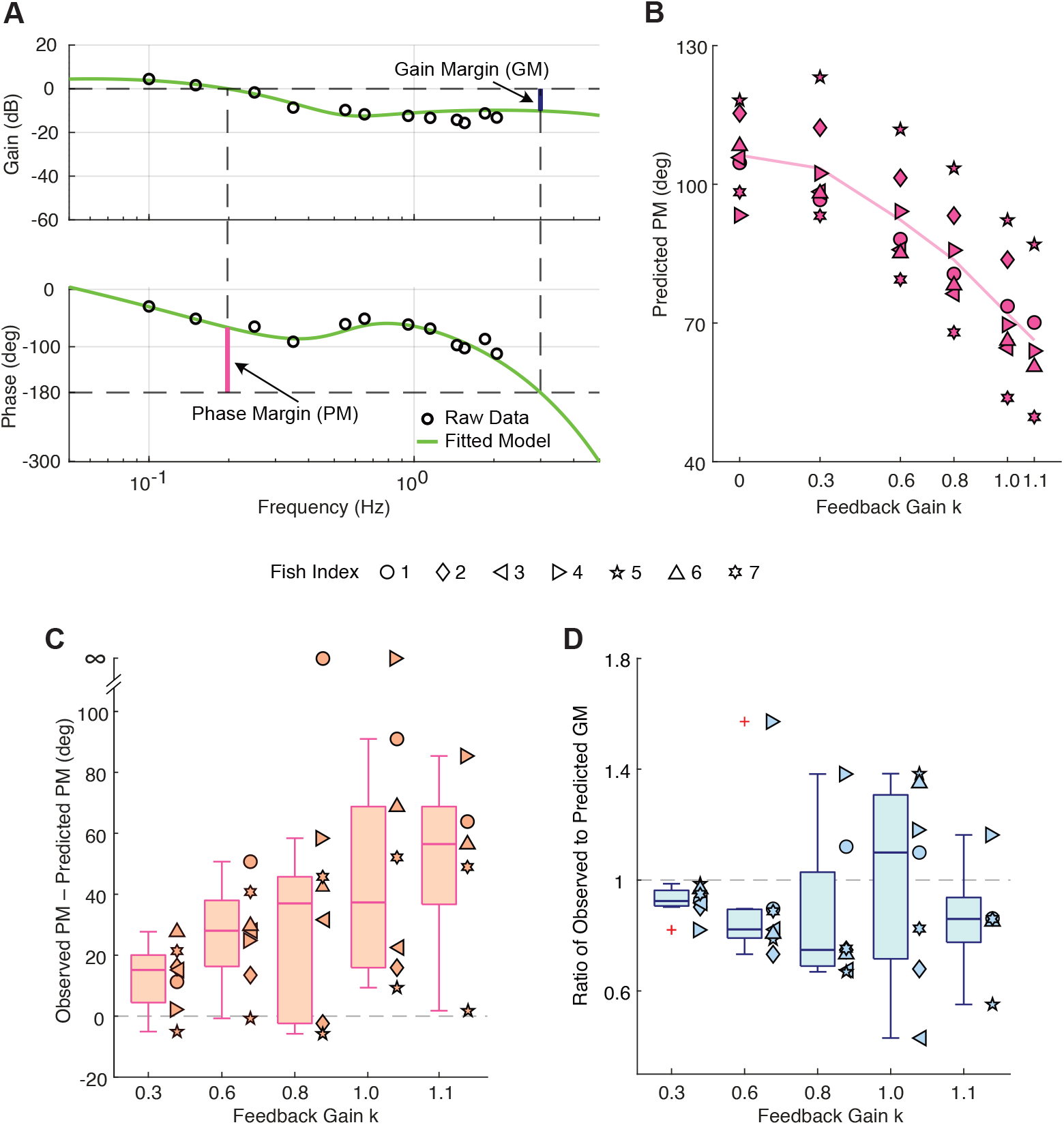
Tuned controller improved phase margin and maintained gain margin, measures of stability robustness. (A) Illustration of the *Phase Margin* (PM) and *Gain Margin* (GM) with raw data (black circles) and fitted model (green line, details in Material and Methods section) from one fish in *k* = 0.8 case. When computing the PM, the frequency where gain crosses 0 dB was within the frequency band we tested, while the frequency where phase crosses 180*^◦^* was beyond 2.05 Hz which therefore required model extrapolation when computing GM. (B) Predicted PM in *k* = 0 (baseline) and all five closed-loop cases from seven fish, computed from the model. As feedback gain increased, the predicted PM markedly dropped, showing that the experimental design destabilized the whole closed-loop system. (C) Box plot of the difference between observed and predicted PM as a function of feedback gain *k* for seven fish (computed from model). The difference is significantly greater than 0*^◦^* in all feedback cases (*p <* 0.05, paired-sample *t* test), indicating that the tuned controller enhanced PM. Note that two cases (fish 1, *k* = 0.8 and fish 4, *k* = 1.0) were excluded from the statistical analysis, because they had infinite observed PM (loop function gain never crossed 0 dB) despite a finite predicted PM. (D) Box plot of the ratio of observed to predicted GM as a function of feedback gain *k* for seven fish (computed from model); a value *>* 1 (or *<* 1) implies the tuned controller increased (or decreased, respectively) GM. The ratio is not consistently above or below 1 as a function of gain, and there is no significant correlation (*p >* 0.1) between high-pass gain *k* and the ratio of observed to predicted PM averaged across seven fish. Therefore, the tuned controller did not significantly enhance (or impair) GM.

For the class of sensorimotor dynamics considered here, these measures of robustness are simple and intuitive: if at any frequency, the loop function *L* reaches the *−*1 point, then at that frequency, the overall negative feedback system experiences *positive* feedback, rendering the closed-loop system *unstable*; further, note that *−*1 point has unity gain (0 dB) and *−*180*^◦^* phase. Thus, if *L* gets close to the *−*1 point, then small changes in the musculoskeletal dynamics could push the system into instability. Thus, the gain margin and phase margin are measures of stability robustness, i.e., measures of the “safety margin” between the observed loop function and the *−*1 point.

If the fish were to have maintained its baseline performance, the PM would have decreased substantially as we increased the artificial feedback gain *k* (Fig. 6B). This predicted PM verified the experimental design, where the goal was to destabilize the system with increasing artificial feedback. However, the difference between observed PM and predicted PM was significantly above 0*^◦^* for all feedback gains (*p <* 0.05, paired-sample *t* test; Fig. 6C)). Moreover, as *k* increased, the difference between observed PM and predicted PM increased (*p <* 0.01, correlation coefficient *ρ* = 0.97 for the average across seven fish). Note that the statistical results of PM did not include data from one fish in feedback case *k* = 0.8 and a different fish in case *k* = 1.0 because both of their observed loop functions did not cross 0 dB, although these data points further support our finding (i.e., infinite observed PM versus finite predicted PM). In summary, fish improved their PM compared to prediction.

For the GM, we observed that as the artificial high-pass feedback gain *k* was increased, the average ratio across seven fish of observed to predicted GM remained near 1 (Fig. 6D). At two gains (0.3 and 0.6), the tuned controller decreased GM when outliers were removed (before removing outliers: *p <* 0.05 for *k* = 0.3, *p >* 0.1 for remaining cases, paired-sample *t* test; after removing outliers: *p <* 0.01 for *k* = 0.3 and *k* = 0.6, *p >* 0.1 for remaining cases, paired-sample *t* test). There was no significant trend as a function of *k* (*p >* 0.1, *ρ* = *−*0.13 before removing outliers and *ρ* = *−*0.0014 after removing outliers).

In total, the fish’s tuned closed-loop system was robust with respect to the artificial feedback, which was designed to render closed-loop system unstable. This result indicates that the tuned controller maintained or improved measures of stability robustness compared to what would be the case if the fish had not tuned its controller.

## 5 Discussion

We examined how the weakly electric fish adapted its controller in the face of destabilizing, closed-loop dynamics. The fish tuned its controller to enhance stability robustness and disturbance rejection, and the learned controller was temporarily retained after the destabilizing feedback was removed, suggesting a cerebellar or cerebellar-like adaptation process. We also found that the fish tuned their controllers in such a way that their stability was *robust*, i.e., small changes in the swimming dynamics or controller parameters were less likely to cause the system to become unstable than had the fish not tuned its controller. In particular, we found that the tuned controller enhanced the phase margin, possibly making the system more robust to the change of illumination or conductivity which might impact sensorimotor delay [36], and thus impact the phase of the loop function.

The tuned controller reduced the impact of disturbances, such as neuromuscular noise and turbulence. In natural biological systems, disturbances can take a variety of forms. For example, humans can robustly reject mechanical disturbances to their hand during reaching [37]. Likewise, when a hawkmoth hovers in a flower’s wake in windy environments, vortices are shed, perturbing the hawkmoth as it feeds from the swaying flower [38]. Future field work could shed light on the relationship between closed-loop disturbance rejection, its sensorimotor adaptation, and the natural statistics of environmental perturbations. This would be similar to the work of Sponberg et al. (2015) [36], which examined the relation between closed-loop tracking performance of hawkmoths and the natural movement frequencies of flowers.

Our present paper focuses on learning of the feedback pathway, however learning can occur in both the feedforward and feedback pathways. Kasuga et al. (2015) [39] discovered that feedforward adaptation and feedback adaptation were separately learned when human participants were trained to reach to two lateral and one central target with mirror-reversed visual feedback. Yang et al. (2021) [28] examined adaptation and *de novo* learning in visuomotor rotation and mirror-reversal tasks, and suggested that there could be two different control pathways, with distinct mechanisms of learning in each. Feedforward control focuses on enhancing tracking and stimulus prediction (inverse internal model), while feedback control focuses on stability and disturbance rejection (forward model). Feedback adaptation involves learning the rich feedback dynamics necessary for stability and performance. Future work could begin to put together these two control elements—feedforward and feedback control—to understand how they are learned in tandem to improve sensorimotor performance in the face of novel dynamics—like riding a bicycle, balancing a pole, and juggling.

While the present paper focuses on the important role that the fish neural controller plays in robust locomotor behavior, it is ultimately the interplay between the mechanical (musculoskeletal) system and neural system that determines stability and performance [15, 18, 40–43]. For example, Sefati et al. (2013) showed that the mutually opposing forces produced by the electric fish ribbon fin enhances both stability and maneuverability, simplifying the role of their neural controllers when maintaining the stability and performance of the overall system [24]. Salem et al. (2022) [9] revealed that flies with damaged wings (i) sacrificed yaw gaze stabilization performance and (ii) actively increased their damping coefficients to maintain stability in yaw gaze stabilization task. The former was from a decrease of closed-loop gain in coupled body dynamics and neural controller of flies and the latter could be due to an increase in wingbeat amplitude and frequency in body dynamics. Full et al. (2002) [44] emphasized that neural control models can be poor and even destabilizing when formulated without understanding the underlying musculoskeletal dynamics. Indeed, it is this adaptive interplay between neural control and muskuloskeletal dynamics that remains the envy of robotics engineers.

## 6 Acknowledgments

We thank Balázs P. Vágvölgyi for modifying the experimental system, Lauren N. Peterson and Huanying Yeh for data collection and analysis, Eric S. Fortune and Sarah L. Poynton for critical feedback on the manuscript, and Christopher S. Yang, Debojyoti Biswas, and Di Cao for helpful discussions. We thank Kyle T. Yoshida, Ismail Uyanik, and Erin E. Sutton for pilot experiments. This work was supported by the Office of Naval Research under grant no. N00014-21-1-2431 (N.J.C).

## 7 Author Contributions

Y.Y., D.G.Y., and N.J.C. co-designed the experiments; Y.Y. and D.G.Y. developed analysis methods; Y.Y. collected, processed and analyzed data; N.J.C. oversaw data analysis, supervised the project, and obtained funding; Y.Y. and N.J.C. co-wrote the original draft.

## 9 Materials and Methods

### 9.1 Subjects

Adult weakly electric glass knifefish *Eigenmannia virescens* (length: 10 – 15 cm) were obtained from commercial vendors and housed following the guidelines [45]. Temperature and conductivity of the water in the experimental aquarium were kept at 76 *^◦^*F – 80 *^◦^*F and 10 *µ*S/cm – 150 *µ*S/cm, respectively. All fishes were transferred to the experimental aquarium 12 – 24 hours before the start of the experiments. The experiments were conducted under an illuminance level of around 80 lux. All the experimental procedures were approved by the Johns Hopkins Animal Care and Use Committee and followed guidelines established by the National Research Council and the Society for Neuroscience.

### 9.2 Experimental Apparatus

The experimental apparatus was modified from the system used by Biswas et al. (2018) [19], which uses a real-time measurement of the fish position, and feeds it back to the input. We modified the system in two ways: (i) the fish position was filtered through a high pass filter before feeding it back to the input, and (ii) the feedback signal was added to a pre-designed sum of sines signal [31]. We refer to our high-pass-filtered feedback as “artificial reafferent feedback.”

### 9.3 Experimental Protocol

Our intention was to drive the entire closed-loop system towards instability by increasing the gain in artificial feedback, and test whether there were aftereffects after the artificial feedback was turned off.

Each experiment comprised three periods: “baseline,” “closed-loop,” and “aftereffect.” Both “baseline” and “aftereffect” periods were open-loop (no high-pass-filter feedback); the “closed-loop” period had the high-pass-filter feedback.

In the “baseline” period, we conducted five 60-second trials, with a 20-second rest period between each trial. The input stimulus in each trial comprised a 10 seconds ramp-up, 40 seconds of sum-of-sines inputs, and 10 seconds ramp-down (see Fig. S1A). Functions are described in section 9.3.1. There were no pre-designed inputs during rest periods.

In the “closed-loop” period, there was a first order high-pass filter in the artificial feedback loop. The high-pass filter gain *k* was increased incrementally: from 0.3, 0.6, 0.8, and 1.0, to 1.1 (Fig. 1C). In each feedback case (*k* = 0.3, 0.6, 0.8, 1.0, 1.1), there were five trials (60 seconds each) with four rest periods (20 seconds each) in between. Critically, the artificial feedback remained enabled during these rest periods.

There were also 20-second rest periods connecting the last trial of each feedback case to the first trial in the next feedback case. During these intertrial rest periods, the feedback gain was linearly increased as a function of time from the last value of *k* to the next value. During all rest periods in the “closed-loop” period, no pre-defined reference inputs were used.

At the start of the “aftereffect” period, the artificial feedback was extinguished. Experiments were conducted based on five time intervals after the end of the closed-loop experiments: “0 to <5 mins after,” “5 to <10 mins after,” “10 to <15 mins after,” “15 to around 22 mins after,” and “approximately one day after.” The input stimulus and the duration of rest periods were the same as those in “baseline” period. Three trials were conducted for “0 to <5 mins after,” “5 to <10 mins after,” and “10 to <15 mins after” cases due to time limits. And five trials were conducted for “15 to around 22 mins after” and “approximately one day after” cases.

#### 9.3.1 Stimulus Design

The pre-designed reference signal had three periods: “ramp-up,” “sum-of-sines,” and “ramp-down,” as illustrated in Fig. S1A. In “ramp-up” and “ramp-down” periods, the designed signal oscillated at 0.45 Hz with modulated amplitude, and was added to the modulated sum of sines signal (Fig. S1A). The sum of 12 single sinusoidal functions, *r*_1_(*t*), is given (in centimeters) as follows:

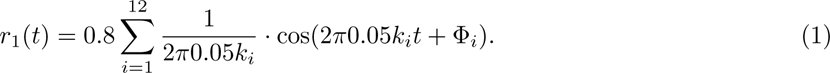

Here, *k_i_* = 2, 3, 5, 7, 11, 13, 19, 23, 29, 31, 37, 41 are prime numbers and the phase of each single sine component Φ*_i_* is randomized. This function creates a pseudo-random, unpredictable stimulus, suitable for system identification [23].

### 9.4 Data Analysis

#### 9.4.1 Online and Offline Tracking

The refuge position and fish position were both measured in real time by a template matching tracker in LabVIEW [19], with a sampling rate of 25 frames per second. The online tracking data was saved in .csv files, and refuge tracking videos from each period of the experiments were also saved as .mp4 files.

To avoid the impact of apparatus latencies on the accuracy of estimating the frequency response of fish in each feedback case, we individually tracked the fish and refuge motion on each video with offline tracking software DeepLabCut [46]. All offline data was finally saved in .csv files with the 2-D position of refuge and fish, together with the tracking likelihood for each frame per body part.

#### 9.4.2 System Identification

After obtaining the refuge and fish position in the time domain, we performed system identification with seven fish. The system identification procedure followed the steps of sum-of-sines based system identification used by Yang (2020) [47]. For each of the three experimental periods (baseline, closed-loop, and aftereffect), we selected the middle 40-second sum-of-sines part of each trial and divided it into two halves to double the replicates of data. We then took the average of each batch of 20-second input and output data in each case (baseline, five feedback cases in “closed-loop” period, and five time intervals in “aftereffect” period) and transformed the averaged data into the frequency domain using the discrete Fourier transform (DFT). We finally used the empirical transfer function estimate (ETFE) to estimate the Frequency Response Function (FRF) for each case. Note that for two fish (fish 1 and 4), due to data quality of “baseline” trials and the similarity of trials in “baseline” and “approximately one day after”, we combined the time domain data in “baseline” and “approximately one day after” cases to estimate the “baseline” FRF.

Trials with tracking loss were discarded. Data from two fish were excluded for *k* = 1.1 due to the numerous trials with tracking loss.

#### 9.4.3 Bootstrapping Analysis of Aftereffect

One of the most important questions of this research was to determine whether the novel closed-loop dynamics had an aftereffect on fish tracking behavior. To answer this question we analyzed online tracking data from six fish, (fish 3 was excluded due to data collection failure). For each 20-second replicate in five cases (“0 to <5 mins after,” “5 to <10 mins after,” “10 to <15 mins after,” “15 to around 22 mins after,” and “approximately one day after”), we excluded those data when there was any tracking loss in any 20-second replicate from any fish and performed bootstrapping (*n*_boot_ = 10, 000) across six fish in the time domain. Then, we estimated the FRFs of 10,000 bootstrapped time domain trials. We computed the mean and standard deviation (SD) of 10,000 bootstrapped FRFs for each replicate. We also followed the same procedure to compute mean and SD of 10,000 bootstrapped FRFs in “baseline” and “closed-loop (*k* = 1.1)” cases, so that we could compare those with bootstrapped FRFs in “aftereffect” cases.

#### 9.4.4 Normalized Error and Averaged Frequency Response Error

To evaluate fish tracking performance, the “normalized error” was computed in the frequency domain by taking the magnitude of the ratio of fish averaged “sensory slip” *E*(*jω*) in each case to reference signal *R*(*jω*) (at frequency components contained by the reference signal). The frequency domain “sensory slip” *E*(*jω*) and the reference signal *R*(*jω*) were computed using DFT. The “sensory slip” *e*(*t*) of fish in the time domain was the subtraction of time domain refuge position *s*(*t*) and output fish position *y*(*t*). From the fish’s “baseline” controller, we predicted the normalized error in each feedback case, and compared that with the observed normalized error from fish tuned controllers. To quantify general impacts of the tuned controller on tracking error, across the frequency range 0.1 *−* 2.05 Hz, we took the integral of the normalized error (using trapezoidal integration over that range), and then divided this integral by the frequency range to obtain the “averaged frequency response error.” Note that although in Figure 4A the *x* and *y* axes are both shown using a logarithmic scale, we computed the intergal of the normalized error in a linear scale.

#### 9.4.5 Stability Robustness Analysis

The two margins of stability (gain margin and phase margin) reflect the stability robustness of the closed-loop system. When the margins of stability decrease, the system gets less robust. To drive the closed-loop system towards instability, the transfer function of the feedback was designed as a high-pass filter in which gain was incrementally increased, causing a decrease of the phase margin. This approach was based on a prediction from baseline, by assuming that the fish’s controller remains the same during the closed-loop experiments.

Here, we computed the loop function

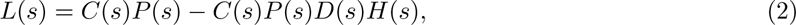

where 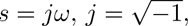

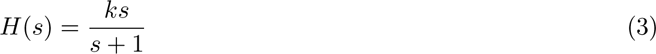

is a first order high-pass filter, and *D*(*s*) is the system delay in the forward loop (see SI Appendix, Supporting Information Text section 10.1.3 for details). The multiplication of transfer function of “controller,” i.e., *C*(*s*), and that of “plant,” i.e., *P* (*s*), could be computed from the estimated fish’s FRFs by opening the fish’s reafferent feedback loop. The transfer function in the artificial feedback was known, so the loop function was then computed, and another array of complex numbers was obtained.

A key question was to determine how the modified controller during closed-loop experiments impacted the stability robustness of the closed-loop system. To address this, we compared the observed gain and phase margins from the fish loop function at each feedback gain case with predicted gain and phase margins.

#### 9.4.6 Bode Sensitivity Integral

The sensitivity function is defined as follows:

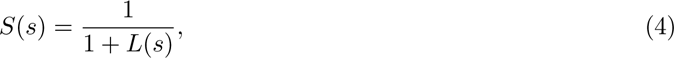

where 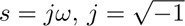. Thus, we could directly compute the sensitivity function from the loop function.

To calculate the Bode sensitivity integral over the frequency range 0.2*π −* 4.1*π* rad/s, we subtracted the positive and negative trapezoidal regions formed by the natural log of the sensitivity function magnitude and the *x* axis (Fig. 5A).

#### 9.4.7 Model Fitting

To find the gain margin which needed data extrapolation, we fitted fish controller times plant *C*(*s*)*P* (*s*) data with models. We fitted data with a second order model (relative degree one: 2 poles, 1 zero with delay)

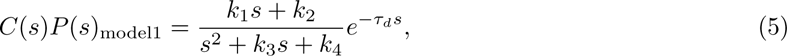

and a third order model (relative degree two: three poles, 1 zero with delay)

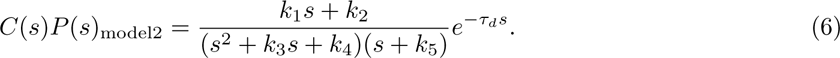

Both models fitted the *C*(*s*)*P* (*s*) data well. The third order model fitted the data slightly better than the second order model as expected (because the former contained one additional parameter); however, the third order model was prone to overfitting. Therefore, for the purpose of analysis, we used the simpler second order model in Eq. (5). The fitted parameters were chosen by the metric defined in SI Appendix, Supporting Information Text section 10.1.2.

#### 9.4.8 Statistics

All the statistical analysis was performed in MATLAB (Mathworks, Natick, Massachusetts, USA). Specifically, we used function corrcoef to test correlation coefficients of data points. We used function ttest to perform pair sampled t-test, and used function mwwtest [48] for the two tailed Mann-Whitney-Wilcoxon (MWW) test. For all tests, the significance level was set to 0.05.

### 9.5 Data Availability

Before final submission, an archived version of the datasets and the analysis code supporting this article will be made available through the Johns Hopkins University Data Archive and will be assigned a permanent digital object identifier (DOI).

## 10 SI Appendix

### 10.1 Supporting Information Text

#### 10.1.1 Conditions of bode sensitivity integral equals zero

Our results introduced a “waterbed effect”, whereby the system suppressed the disturbances at some frequencies at the cost of enhancing disturbances at other frequencies, because the Bode sensitivity integral equals zero. This result requires that the *loop function L* of the feedback system satisfies two conditions: 1) has no unstable poles, and 2) has at least two more poles than zeros.

The electric fish plant dynamics *P* (*s*) have been well approximated by Sefati et al. (2013) and Uyanik et al. (2020) [15, 24], using a stable second order model with no zeros. Assuming the fish controller *C*(*s*) is causal, and thus is proper (degree of denominator *≥* that of numerator), it is easy to show that *loop function L* = *CP − CPDH* satisfies the two conditions above for the Bode sensitivity integral to be 0. We could clearly observe a “waterbed effect” (Fig. S3), which shows both raw data and the fitted sensitivity function in *k* = 1.0 case, with the relative degree two model in Eq (6) in the frequency range of 0 *−* 2*π ×* 25 rad/s.

#### 10.1.2 Model fitting and “best” parameter selection

To select the “best” parameter model, we followed three steps.

**Step 1: Define the “fitting error.”**

The frequency response of each feedback case was represented as a vector of 12 complex numbers:

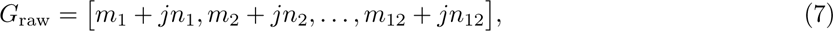

one for each frequencies that was contained in the reference sum-of-sines stimulus [23, 49], where 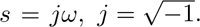.

Similarly, we obtained frequency response functions at these 12 frequencies directly from the fitted model with *s* = *jω*, noted as:

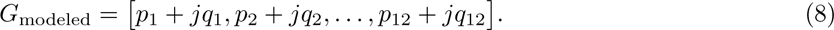

The fitting error is defined as the mean-squared error between *G*_raw_ and *G*_modeled_, i.e.,

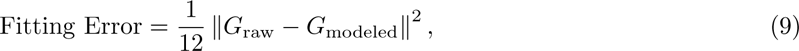

where *||·||* is the 2-norm of vector.

**Step 2: Use MATLAB function to obtain sets of optimal parameters in candidate model that minimize the fitting error corresponding to different initial conditions.**

As an example, the second order model (relative degree one: 2 poles, 1 zero with delay) has five parameters: *k*_1_, *k*_2_, *k*_3_, *k*_4_, and *τ_d_*. We randomly generated 500 sets of initial values for each of those five parameters. Then we used MATLAB function “fminseasrch” to find 500 sets of optimal parameters [*k*_1_, *k*_2_, *k*_3_, *k*_4_, *τ_d_*] that made the fitting error reach the local minimum from each initial condition.

**Step 3: Select the least fitting error among sets of optimal parameters.**

Finally, we selected a set of parameters, among all 500 sets of optimal parameters corresponding to each initial condition, that gave the least fitting error.

#### 10.1.3 System Delay Analysis

The latencies in the experimental setup can impact the accuracy of estimated fish’s FRF in system identification, and this needs to be taken into consideration when analyzing our data. In our system, the delay was mainly the latencies between reading the reference files from the PC and trigger of the linear actuator to control the refuge movement. There were only negligible delays caused by other factors such as asynchronously saving the videos and .csv files, and the artificial feedback processing delay from FPGA.

In control theory, delay can easily be modeled by the transfer function *e^−τs^*, where *τ* is the time constant. Here we denote the system delay as *D*(*s*). To generate the offline refuge movement data, we processed recorded experimental videos with the tracking software DeepLabCut [46], and estimated the delay in the frequency domain by taking the ratio of the Fourier domain offline refuge trajectories, to the Fourier domain online refuge trajectories saved from the experimental setup.

#### 10.1.4 Disturbances in Fish Refuge Tracking Task

Sefati et al. (2013) [24] explained that the forward-backward movement of *E. virescens* relies on mutually opposing thrust forces generated by two inward, counter-propagating waves of its ribbon fin. The change of thrust forces is linearly controlled by the fish adjusting where these two waves meet (i.e., “nodal point” position). The authors also found a damping force, opposing the direction of the velocity of the fish during tracking. That damping force arose from asymmetries in net velocities of the counter-propagating waves caused by body forward-backwards velocity [24].

Based on these findings, Sefati et al. (2013) [24] constructed a simple, lumped-parameter, task level, dynamical model for the “plant” of the fish:

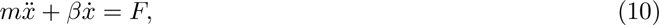

where *m* is the mass of fish, *β* is the damping constant, *F* is the net thrust force generated by nodal point movement of ribbon fin, and *x*, *x*, *x* are the position, velocity, and acceleration of fish in longitudinal direction, respectively. Uyanik et al. (2020) [15] quantified the nodal point position *u* (proportional to *F* in Eq (10), with a nodal shift gain *κ*) as the fish’s motor output (input of the plant) and longitudinal position of the fish *x* as system output, i.e.,

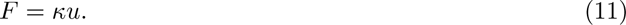

From control theory [32], we know that disturbance of a control system is added to the control input (i.e. motor output in biological system [50]), and the addition becomes input of the plant. When the fish moves back and forth to track the refuge, it causes water turbulence in the tank. That could generate a disturbance force *F_t_* impacting the longitudinal net force in fish tracking. Besides, the motor command noise *d_m_* could directly act on adjusting the nodal point position *u*. Thus, we have

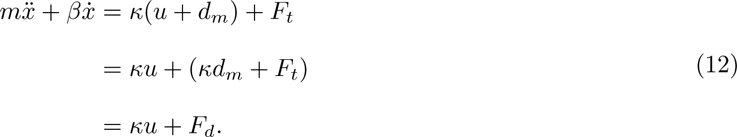

Here, *F_d_* is the combined “disturbance” from turbulence forces and motor command noise, and does not impact the structure of the plant model.

---

**Figure S1.**
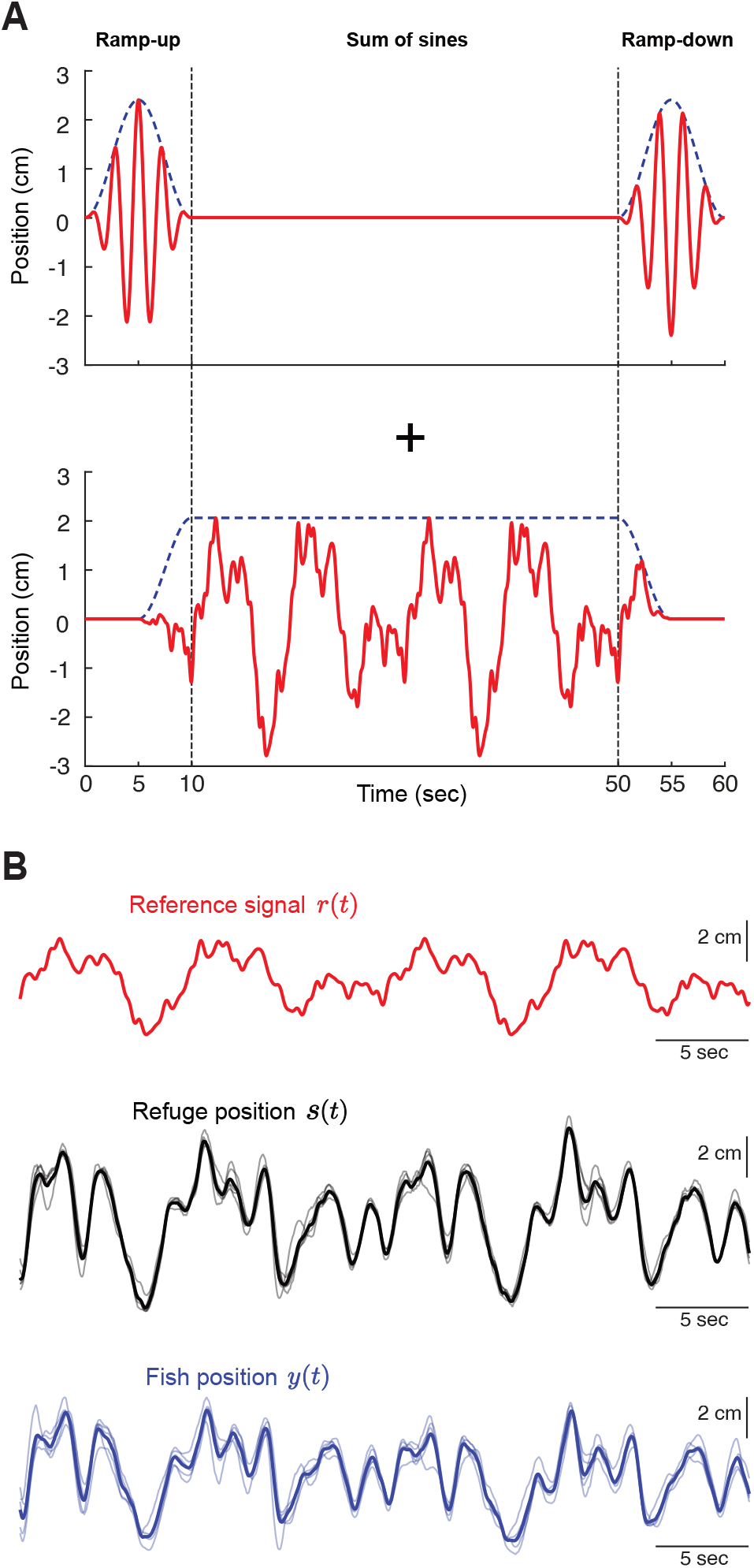
(A) Pre-designed reference signal *r*(*t*). In each 60-second trial, the first and last 10 seconds were “ramp-up” and “ramp-down” periods, in which the pre-designed signal oscillated at 0.45 Hz with modulation (upper part) and was added to modulated sum of sines signal (lower part). The 10 − 50 second was the “sum-of-sines” period, in which the designed signal followed summation of sinusoids in Eq (1). Details are in Materials and Methods Section 9.3.1. (B) Time domain offline data in “sum-of-sines” period (40 seconds) from one fish in the closed-loop *k* = 1.1 case. Red: reference signal *r*(*t*); Black thick: averaged refuge position *s*(*t*) across *n* = 5 trials; Black thin: refuge position in each individual trial; Blue thick: averaged fish position *y*(*t*) across *n* = 5 trials; Blue thin: fish position in each individual trial.

**Figure S2.**
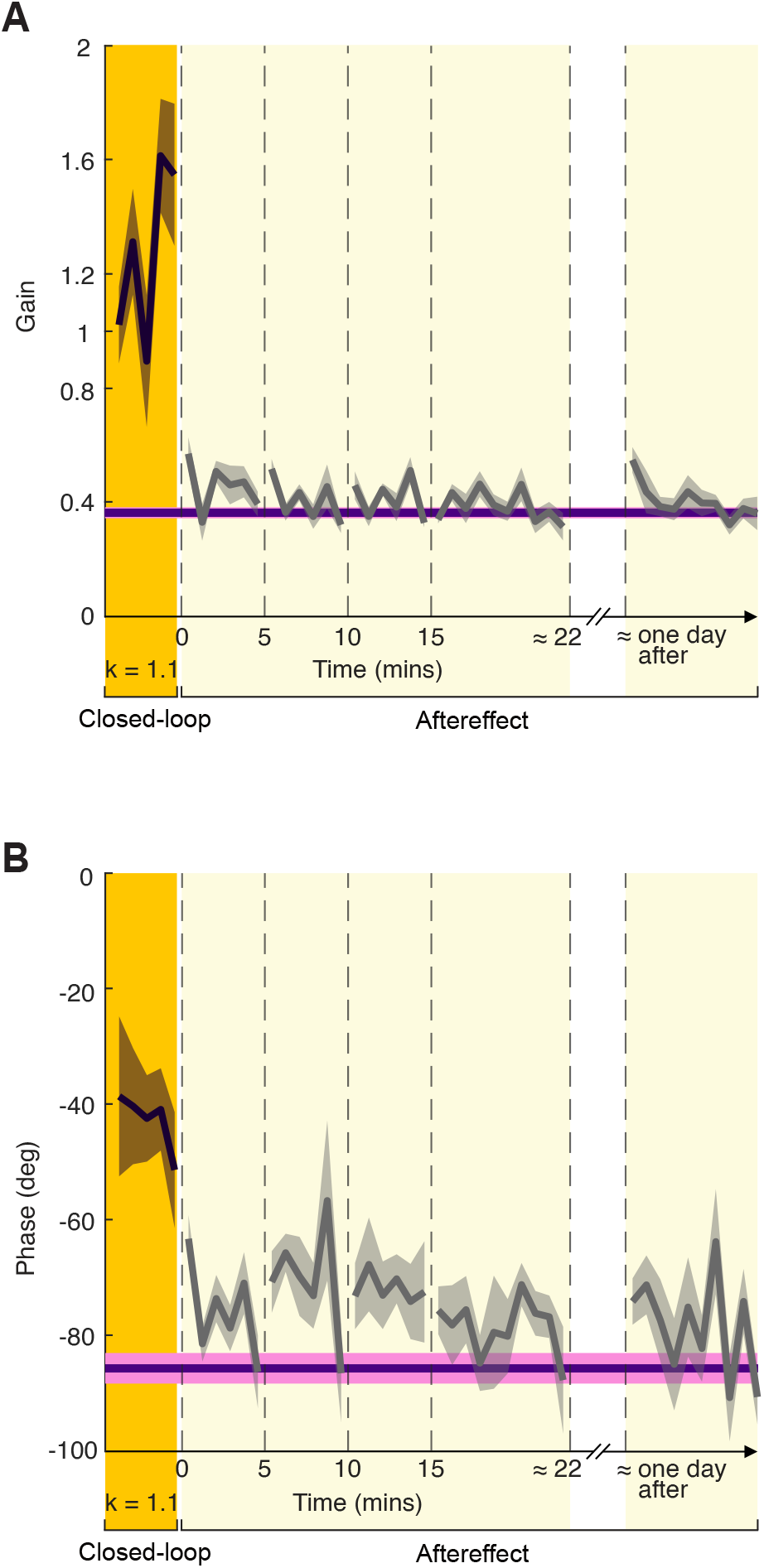
Fish FRF gain at 1.15 Hz(A) and phase at 0.65 Hz(B) from late learning “*k* = 1.1” (when *t <* 0), “0 to <5 mins after,” “5 to <10 mins after,” “10 to <15 mins after,” “15 to around 22 mins after,” and “approximately one day after” cases, by bootstrapping online data (10,000 times) across six fish. The solid gray lines are mean, and the shaded regions are standard deviations (SDs). The solid purple line is the mean of the bootstrapped data in “baseline,” and the adjacent pink shaded region is SD.

**Figure S3.**
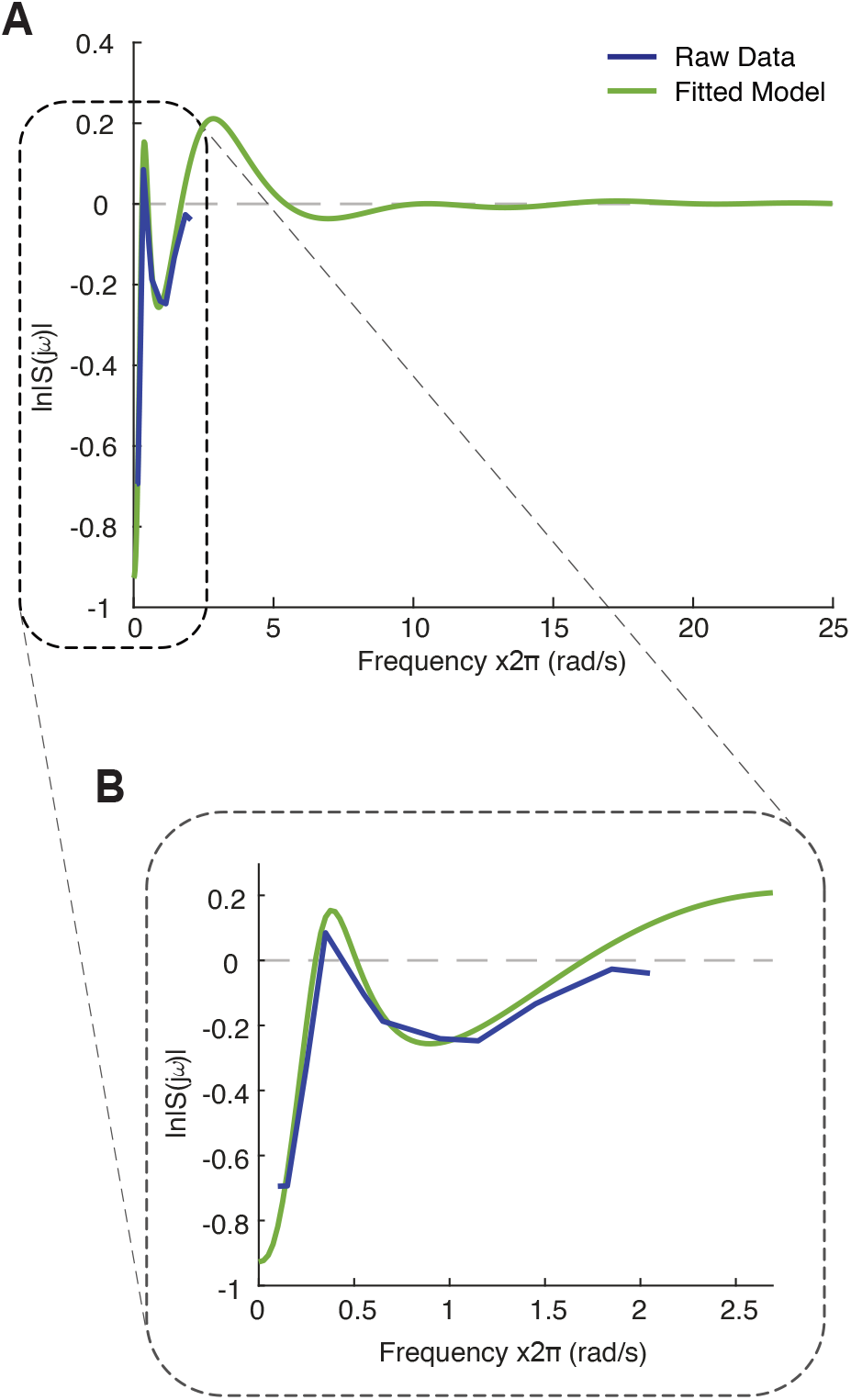
(A) Fitted Sensitivity function in *k* = 1.0 case from one fish (green) with frequency range of 0 2*π* 25 rad/s and raw observed data (blue). A distinct peak above zero indicates that the tuned controller of the fish reduced its sensitivity to low frequency disturbances (*<* 2.05 Hz), possibly as a cost of sacrificing the sensitivity to frequency band 2*π* 1.7 2*π* 5 rad/s (mostly of which was beyond our experimental design). (B) Enlarged version of portion of (A) to show the frequency range in our experiment.

**Figure S4.**
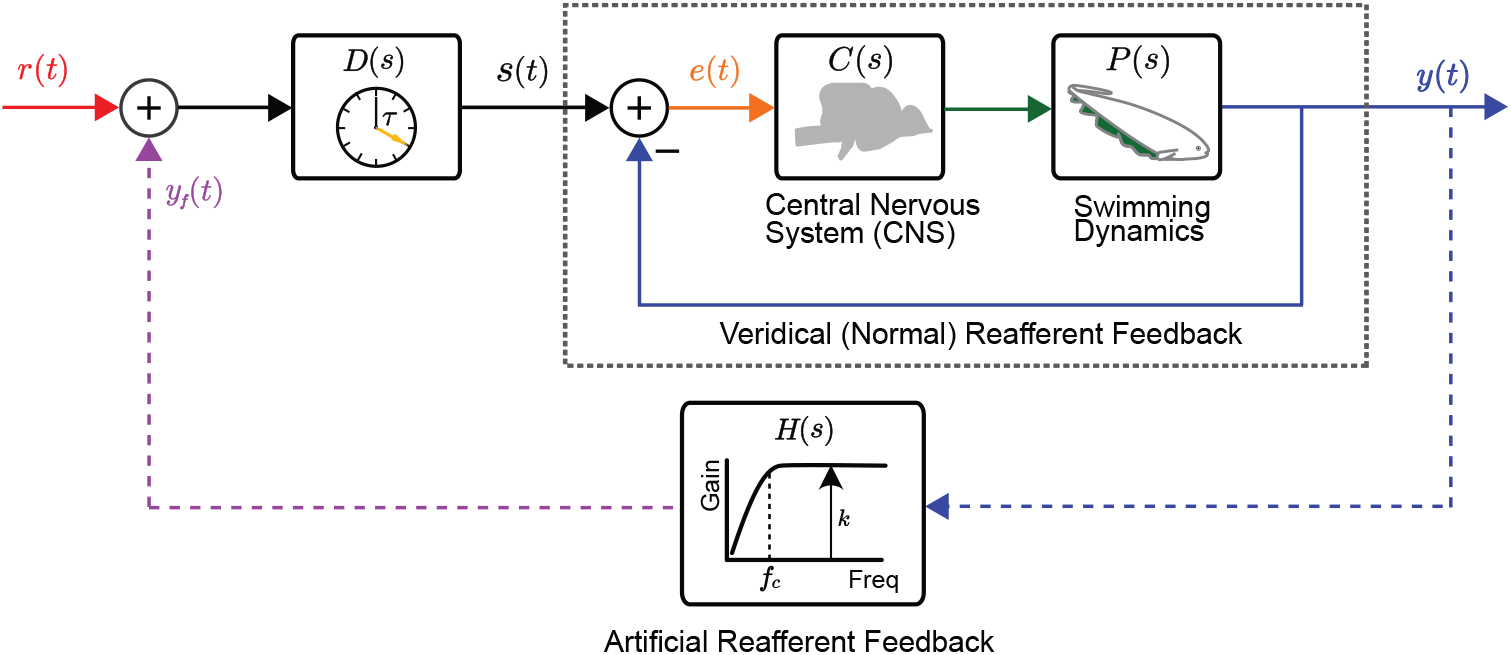
Block Diagram of the fish refuge tracking system with setup delay. The setup delay block *D*(*s*) is at the forward loop. Identification of setup delay helped accurately estimate the sensitivity function.

